# MDRepo – an open environment for data warehousing and knowledge discovery from molecular dynamics simulations

**DOI:** 10.1101/2024.07.11.602903

**Authors:** Amitava Roy, Ethan Ward, Illyoung Choi, Michele Cosi, Tony Edgin, Travis S. Hughes, Md. Shafayet Islam, Asif M. Khan, Aakash Kolekar, Mariah Rayl, Isaac Robinson, Paul Sarando, Edwin Skidmore, Tyson L. Swetnam, Mariah Wall, Zhuoyun Xu, Michelle L. Yung, Nirav Merchant, Travis J. Wheeler

## Abstract

**Background:** Molecular Dynamics (MD) simulation of biomolecules provides important insights into conformational changes and dynamic behavior, revealing critical information about folding and interactions with other molecules. This enables advances in drug discovery and the design of therapeutic interventions. The collection of simulations stored in computers across the world holds immense potential to serve as training data for future Machine Learning models that will transform the prediction of structure, dynamics, drug interactions, and more.

**A need:** Ideally, there should exist an open access repository that enables scientists to submit and store their MD simulations of proteins and protein-drug interactions, and to find, retrieve, analyze, and visualize simulations produced by others. However, despite the ubiquity of MD simulation in structural biology, no such repository exists; as a result, simulations are instead stored in scattered locations without uniform metadata or access protocols.

**A solution:** Here, we introduce MDRepo, a robust infrastructure that supports a relatively simple process for standardized community contribution of simulations, activates common downstream analyses on stored data, and enables search, retrieval, and visualization of contributed data. MDRepo is built on top of the open-source CyVerse research cyberinfrastructure, and is capable of storing petabytes of simulations, while providing high bandwidth upload and download capabilities and laying a foundation for cloud-based access to its stored data.

## Introduction

In Molecular Dynamics (MD) simulation, the movement and interactions of one or more molecules is estimated over time by calculating the force on every atom at discreet time steps on the order of femtoseconds. MD simulation of the fluctuation of a protein molecule with several thousand atoms is commonly captured over time scales of nanoseconds to microseconds, enabling exploration of molecular interactions at spatial and temporal scales that are difficult to observe experimentally (1). The primary products of MD simulations are coordinates over time, known as trajectories, saved at a userdefined frequency (often pico- to nanoseconds). The resulting size of the files capturing simulated atomic trajectories is on the order of many gigabytes. A wide variety of post-simulation analyses are performed by researchers, ranging from quality control, to free energy estimates, to measurements of molecular mobility.

It is common in most data-intensive areas of biological science that primary data is captured in open repositories, made publicly available when associated research is published. This has been codified in the data management and sharing policies of most journals and large research funding organizations. For example, is now a common funding agency mandate that all generated data should adhere to the FAIR guiding principles for scientific data management and stewardship (i.e., ensuring that data are Findable, Accessible, Interoperable, and Reusable (2)).

This guiding principle has driven the creation of invaluable centralized open-access repositories for large-scale data across bioinformatics. Notable examples include archives of sequence and functional information for proteins (3) and DNA (4, 5), a gene expression atlas (6), a data bank for protein structures (7), and databases for classifications of protein families (8) and structural domains (9). These repositories are characterized by for their support for scalable expansion, performant search and retrieval, structured metadata, and open access nature; most are designed to grow by accepting data contributions from researchers across the globe.

Despite the trend for open access repositories, there is no equivalent option for creators of MD simulations. A few special-purpose databases do exist (e.g. (10–16)), but none are designed to scale to meet community needs. Because no adequate repository currently exists, the existing worldwide collection of protein MD simulations, reaching well into the petabytes in scale, is stored in a highly fragmented landscape. Researchers fulfilling the expectation that their data is made publicly available are forced to either host their own web server or resort to sharing simulation data via one of several unstructured, general-purpose open repositories (e.g. Zenodo (17), OSF (18), FigShare (19)). Meanwhile, the large majority of MD data are stored on private computers with no public access capability.

The lost opportunities resulting from the current fragmented data landscape can hardly be overstated. One obvious consequence is that a researcher who might benefit from a collection of previously-performed simulations is likely to be unaware of their existence, and will therefore either repeat the expensive calculations or proceed with no such simulation data. Arguably more important is the lost potential to use the large collection of existing simulations to train systems (especially machine learning models) for a variety of analytical problems that would benefit from a nuanced understanding of the diversity of dynamics of molecular systems. One compelling example of the role that a large collection of MD simulations could play in training of machine learning models is in the context of rapid computational prediction of drug binding affinity and dynamics. Modern computational protein-drug affinity estimation methods demonstrate limited general predictive power (20, 21); the failure of these models limits their utility in drug development, and stems in great part from insufficient volume of training data (22, 23) and lack of representation of structural variability (24). A large and diverse data set of MD simulations will serve as the launch pad for future machine learning methods in protein-drug affinity prediction (25), just as large-scale protein structure databases provided the necessary training data for transformative deep learning methods for structural prediction (26–28).

It is remarkable that an open repository for protein/drug MD simulations does not currently exist, considering the extensive use of such simulations in research labs around the world, the large computational burden of individual simulation runs, the reusability of resulting data, the increasing emphasis on FAIR data management, and the high value of such data for training machine learning tools to perform a broad spectrum of related analyses. As evidenced by the many existing databases, there is no shortage of interest in creating such a repository, so we suspect that the lack of a general repository is primarily due to the challenges of scale. Considering both published and unpublished simulations, largescale projects, and individual research efforts, it seems likely that several million protein MD simulations have been performed over the decades. Therefore, the total size of existing MD simulation data must range in the many petabytes in size. This data scale places extreme demands on infrastructure, both for storage and the data transmission required to enable convenient access to multiple simulations. These demands necessitate a hardware and system architecture that are generally beyond the scope of a single research group.

Here, we introduce a new service designed to fill this void. *MDRepo* is an open repository that is designed to support community contribution, large-scale retrieval, visualization, and cloud-backed analysis of biomolecule MD simulations.

It is designed to provide a home for the millions of simulations accumulated over decades of research effort, with an expected eventual scale of 10s of petabytes. Storage of simulated trajectories is intended to reduce redundant research efforts, improve reproducibility, and enable new discoveries and modeling techniques. In the initial release, *MDRepo* is built to accommodate protein simulations (with or without ligands); it will soon expand to capture simulations of all biomolecules. Data stored in *MDRepo* are released under the open Creative Commons Attribution 4.0 International License (https://creativecommons.org/licenses/by/4.0/), ensuring unfettered use and distribution of its simulations. In the following sections, we introduce the *MDRepo* user interface and describe its underlying architecture.

## Website and User Interface

Researchers will interact with *MDRepo* primarily through its website. A site user can explore stored simulations, its metadata, and any available results of downstream analyses. They can also identify simulations matching particular search constraints and manage data movement (contribution and download) of any number of simulations. Each simulation is stored as a separate entry, with standardized metadata captured for each. Each entry is assigned a unique and persistent accession number.

### Data Exploration page

The common page for searching and exploring *MDRepo* data is the Explore page. This page, as seen in Figure 1, presents a list of all simulations in the database, and can be sorted and filtered to meet user requirements. Search fields include the simulation “Description”, “Biomolecules” and “Ligands” associated with the simulation, the “Protein sequence”, and the “Software” used to create the simulation (some fields are hidden from view in the screen capture). Fields can be dynamically added and removed from user view. Results are paginated with user-selected page length (default is 10 simulations per page).

**Fig. 1.**
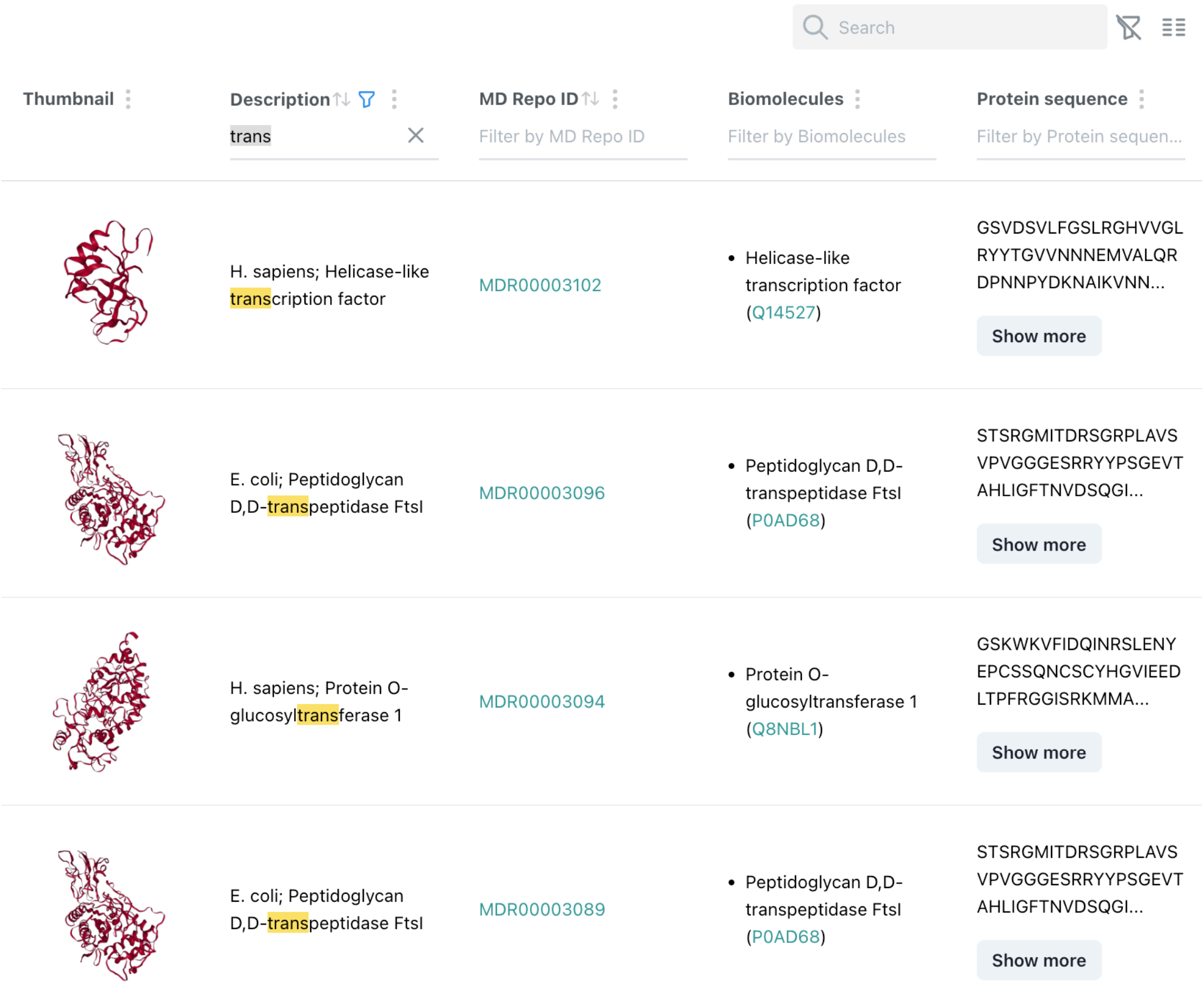
*MDRepo* Explore page, showing a table containing information about some simulations. In this case, the user has filtered for searches matching the partial term “trans”, and a few of the matching results are shown.

### Simulation Detail page

A site user may click on an entry in the Explore page, and navigate to the Simulation Detail page (Figure 2) for a selected simulation. This resulting page provides additional simulation properties, such as duration and time steps, RMSD / RMSF values, simulation software version and parameters, and (where applicable) a linkout to the original website source of the simulation. The Simulation Detail page also contains a visualization of the trajectory as rendered with the NGL viewer, and provides access to download all files associated with the simulation.

**Fig. 2.**
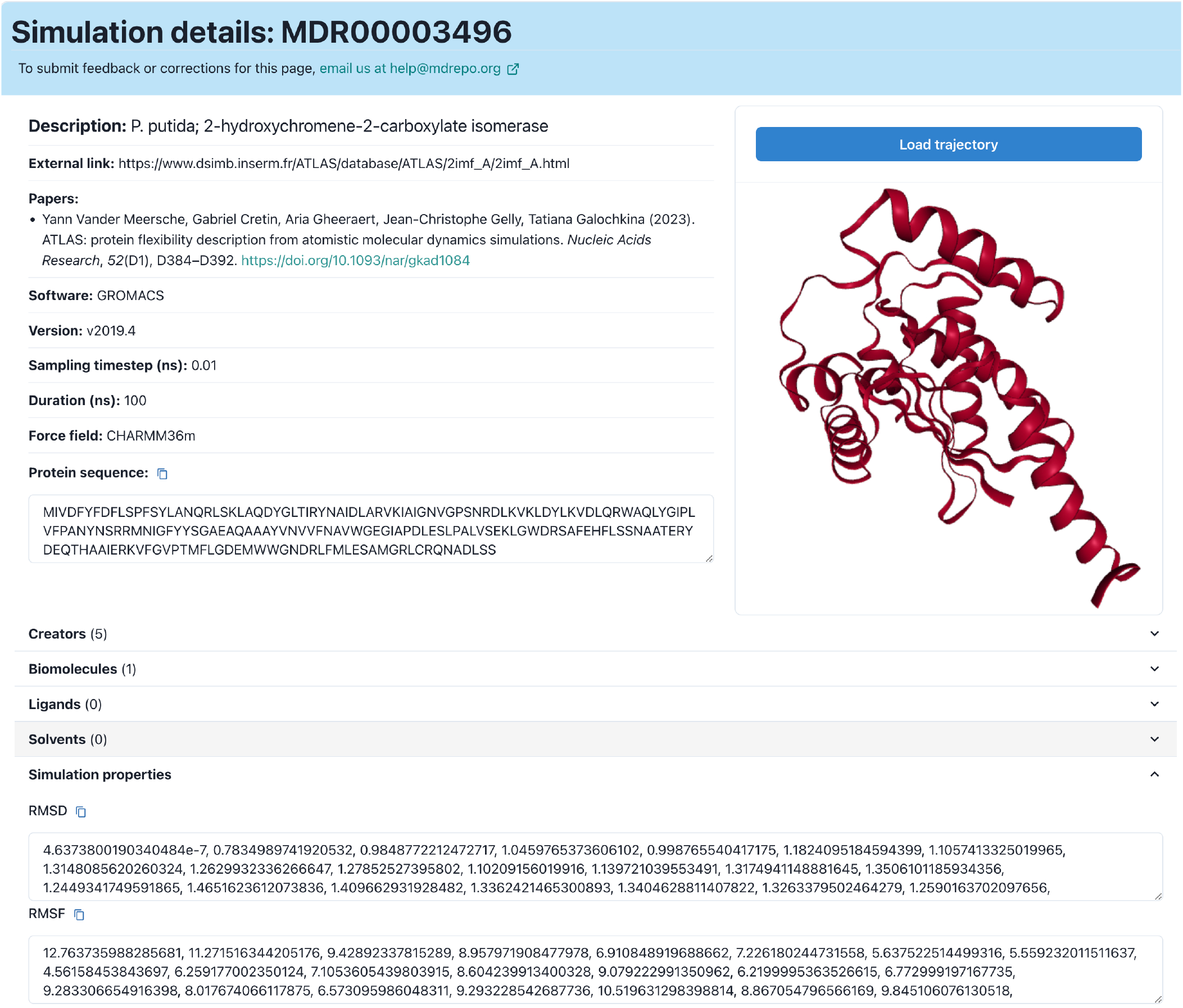
*MDRepo* Simulation Detail sample page, showing the organization of metadata captured and presented for an individual simulation.

#### Data download

Data for a single simulation, including the files associated with the simulation, can be downloaded from the Simulation Detail page, using the list of files presented at the bottom of the page (Figure 3). The selected files will be compressed into a “.zip” file and downloaded through the browser.

**Fig. 3.**
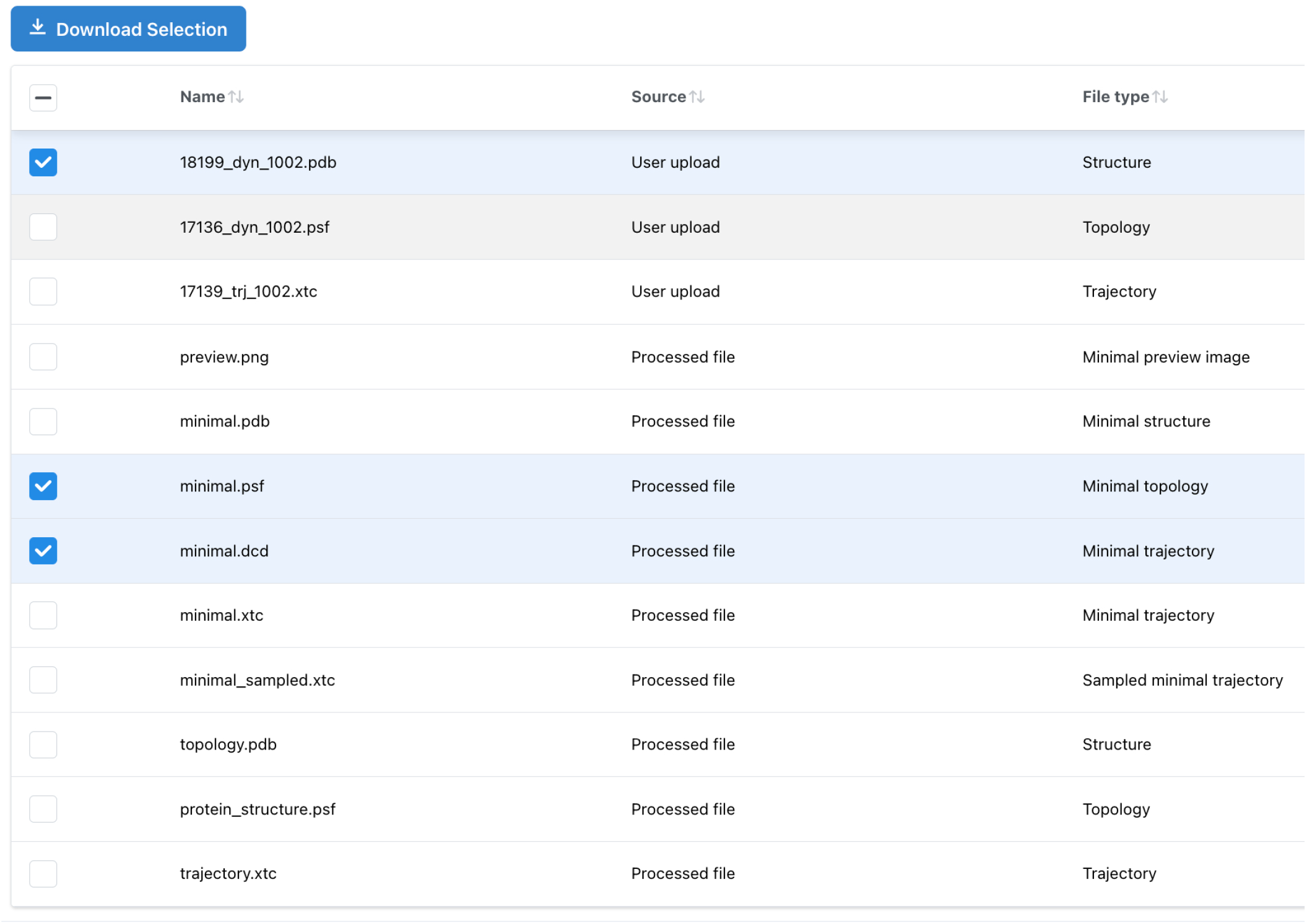
On the *MDRepo* simulation detail page, a user may select and download one or more files associated with a simulation.This figure shows the collection of files available for simulation MDR00001111.

The *MDRepo* system also supports download of multiple simulations at the same time. Rather than downloading batch simulations through the browser, *MDRepo* is designed to support high throughput and fault-tolerant download directly to a user-side server, such as an HPC resource, where there is expected to be both sufficient storage to hold the requested data and sufficient computational power to perform analyses on the downloaded simulations. A user can download many simulations by first selecting the desired entries from the Explore page, then clicking the “Download Selection” button. This causes the backend to generate a download token that contains access information about the simulations to be downloaded. The user must install the *MDRepo* commandline tool (`mdrepo`, https://github.com/MD-Repo/md-repo-cli/) on the recipient server, run the command ‘`mdrepo get`’ as instructed (Figure 4), and supply the token provided as a result of choosing “Download Selection”. Note that the Download Selection button for multiple downloads is only available for users who have signed in with their ORCID iD. This limitation is taken to limit risk of denial of service attack.

**Fig. 4.**
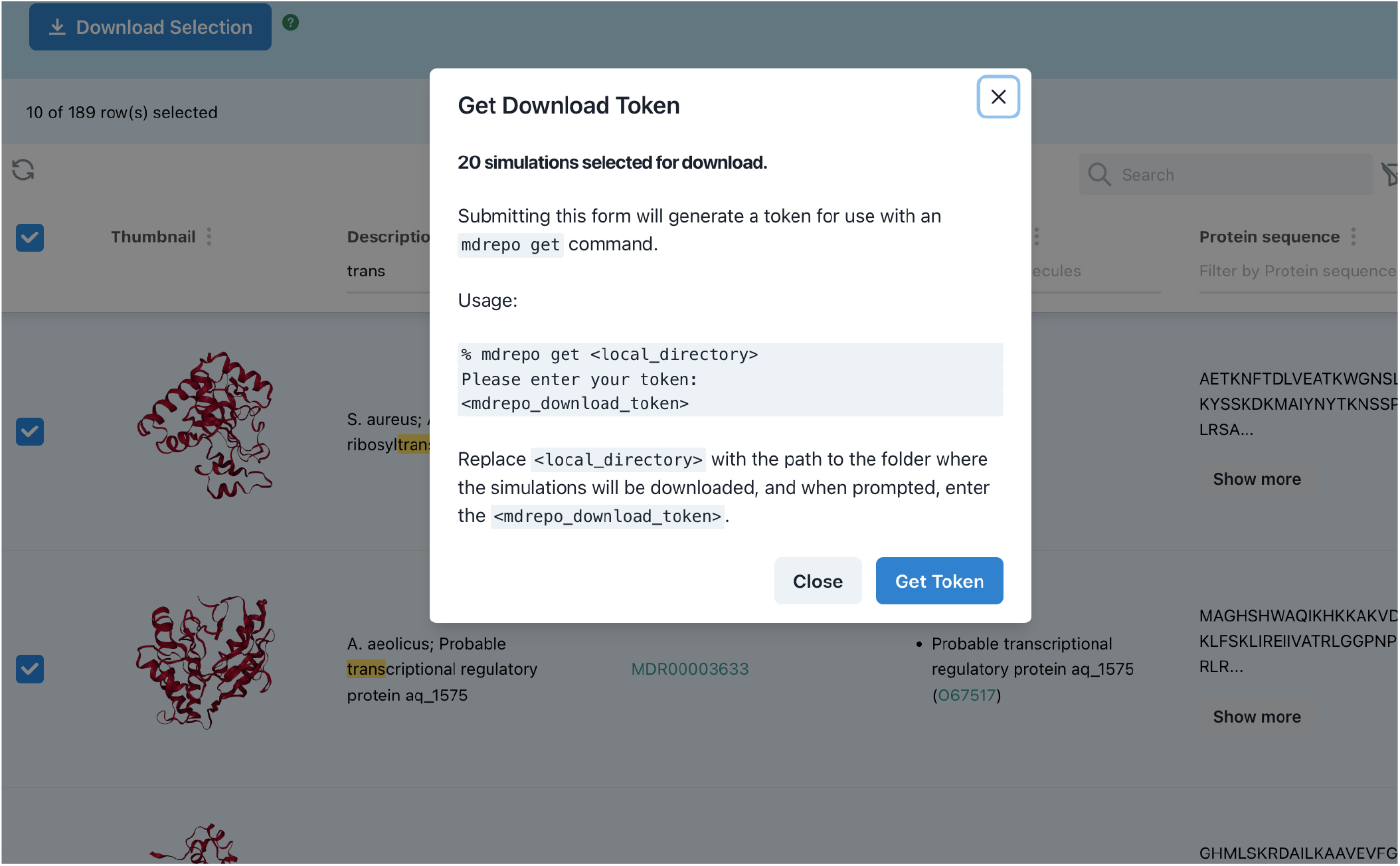
After selecting a batch of simulations to be downloaded, the user clicks the Download Selection button, receives instructions for running the ‘`mdrepo get`’ command, and is provided a token to support download on the recipient computer. For batch download, the computer is expected to be a server, not the system from which the user accessed MDRepo.

### Data submission

*MDRepo* allows contributions from authenticated users. A contributor must create a metadata file for each simulation that they wish to upload, and organize their simulation files in a specific manner. They can then use the `mdrepo` command-line tool to upload their simulation files.

A submission directory is a single directory containing a set of subdirectories, with one subdirectory for each simulation to be uploaded. Each simulation subdirectory contains one trajectory file, one structure / coordinate file, one topology file (.psf for CHARMM, NAMD, XPLOR, .top, .itp, .tpr for GROMACS, .prmtop for AMBER etc.), the simulation metadata file, and any additional files produced with the simulation.

The metadata file for each Simulation can be generated manually (optionally with assistance from the help page provided on the MDRepo website: https://mdrepo.org/metadata), or with a contributor-created script that converts user-structured data into the precise format expected by *MDRepo*. The required metadata includes information such as simulation descriptions, protein/ligand information, the software used to produce the simulation, published papers, contributor details, and information about the files to be uploaded.

Contributors must have the `mdrepo` command-line tool installed on the system containing the simulation directories in order to perform the upload. They may then click the “Contribute” button on any page of the *MDRepo* site, and choose “Get upload tokens”. A cryptographic token is then created, which ensures that the account that generated the token is associated with the person that submits the simulation files from their computer. Simulation upload progress or errors can be tracked on the upload logs page (Figure 5).

**Fig. 5.**
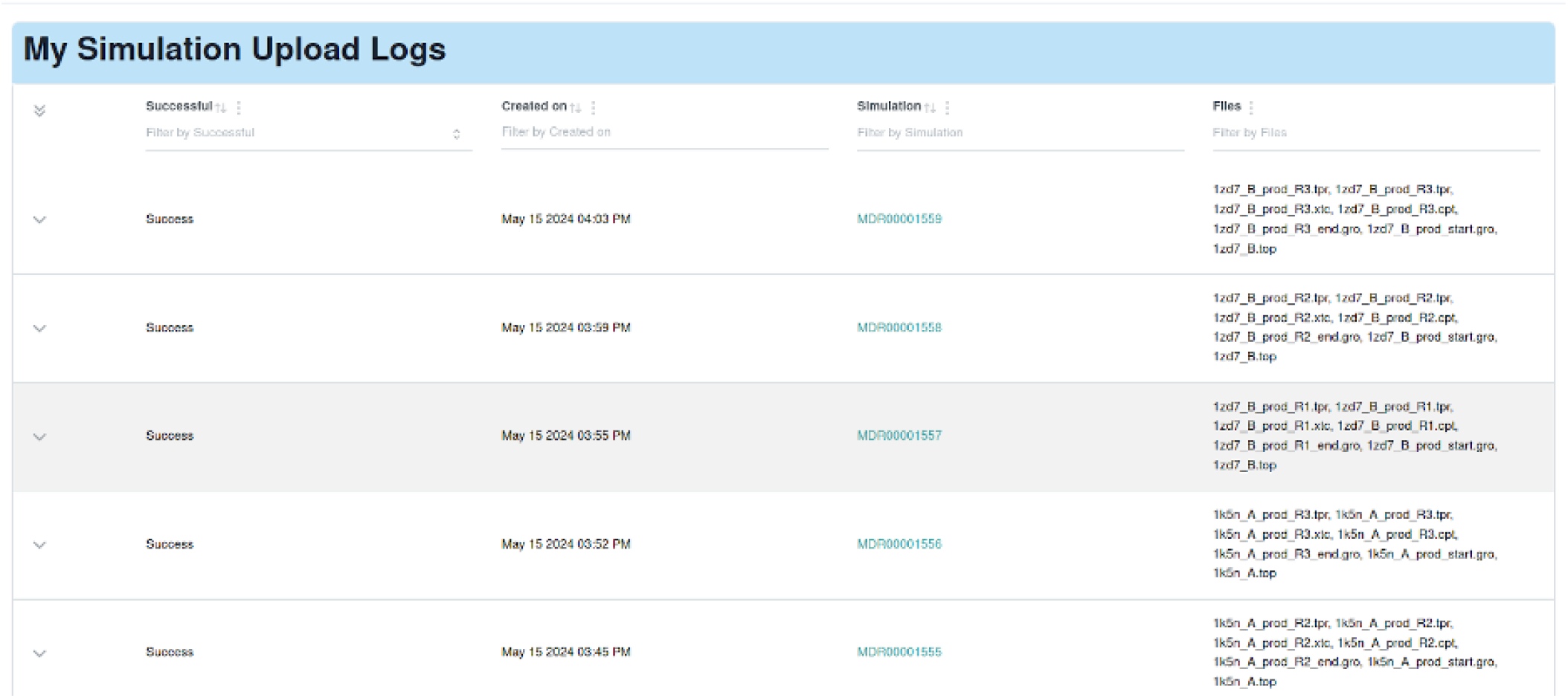
Once a user has contributed one or more simulations, they can view their upload log. The log contains information about ongoing and past contributions.

## Methods

### System Architecture

Users of *MDRepo* will primarily interact with the website (https://mdrepo.org/), where they can explore existing simulations, request batch downloads from the data store, and initiate data contributions. For contributions and large-scale retrievals, the user manages data transfer with our `mdrepo` command-line tool, which controls data upload/download in a high-throughput and fault-tolerant manner.

All functionality rests on the foundation provided by the open-source CyVerse (29) research cyber-infrastructure. *MDRepo* is a cloud native platform deployed onto Kubernetes, a container orchestration engine. This enables MDRepo to scale specific services, such as the web application, in response to increasing connections, cpu load, or RAM utilization. In addition, Kubernetes provides facilities for high availability and load balancing of containerized services. Upgrades can be seamlessly deployed using controlled rollout process. All these features provide for a robust *MDRepo* platform that can scale and grow as the number of users and data grows.

#### Website

The MDRepo website is hosted on a set of virtual machines (VMs) within at the JetStream2 cloud computing environment (30). The front end provides an interactive user experience based on the React (31) and Next.js (32) frameworks, supplemented with Chakra UI (33) components. Visualization of simulations is performed using the NGL viewer (34). Website backend operations are handled by the Django web framework (35), with database operations supported by PostgreSQL (36). *MDRepo* only supports requests for batch upload or download from site users who have authenticated using a valid ORCID account (37) to avoid site vandalism and denial of service attacks; general exploration and single-simulation downloads do not require authentication.

#### Data storage

*MDRepo* content is stored in one of two ways, with content storage divided in a way that can accommodate peta-scale simulation data while providing for a fast and interactive website. (1) Metadata about both simulations and users is captured in a site-specific PostgreSQL relational database hosted on a VM co-located with the primary webserver. (2) All large primary data files (such as trajectory and topology files) are stored in CyVerse Data Store. CyVerse Data Store is managed across two sites: the University of Arizona (UArizona) and the Texas Advanced Computing Center (TACC), and is built on top of the Integrated Rule Oriented Data System (iRODS) (38), a federated data grid and data management system with high throughput data handling capabilities. Data managed through iRODS benefits from the underlying metadata driven rules and policies that afford fine grained access control, along with automation through its message bus architecture and subscription based event monitoring that allows integration with external systems. Data stored in CyVerse Data Store is replicated between UArizona and TACC to ensure data availability, reliability, and resilience.

#### Data contribution and download

We expect that most data uploads will be performed from computer servers where simulations were performed, rather than from researchers’ laptops. Similarly, we expect that downloads involving multiple simulations will generally aim to gather data to a user side server where large-scale analyses can be performed. We have developed a command-line tool to meet these needs, `mdrepo` (https://github.com/MD-Repo/md-repo-cli), written in the Go programming language. The user first requests an MDRepo token from its website (for either contribution or download) to initiate data transfer, then provides that token to the commandline tool for authenticated data transfer.

A path to a directory is provided to `mdrepo` for upload. The directory may contain multiple subdirectories – each is treated as a distinct simulation for submission, and must contain a topology file, a trajectory file, and a metadata file containing information specifically describing the simulation (see https://mdrepo.org/metadata). In the case of download, sub-directories are added to the provided path, one for each requested simulation.

#### Post-upload processing pipeline

Upon completion of simulation upload to the *iRODS* landing directory, a *postprocess* event is initiated in the *MDRepo* backend. This event validates each submitted trajectory, performs a few standard analyses, and loads information from the simulation metadata file into the webserver’s database. File upload status is monitored by the CyVerse Datawatch (https://gitlab.com/cyverse/datawatch) system; after all files in a simulation directory have completed transfer, Datawatch makes a post request to the Django backend, where Django Q2 (https://django-q2.readthedocs.io/en/master/) manages the task queue for the entire submission process.

Data formatting consists of the following steps:

- Verify that the metadata file is valid (correct format, all required fields are present and meet the field requirements).
- Confirm that files specified in the metadata file exist in the upload.
- Ensure that the submission is not a duplicate (based on the file hashes of the uploaded files).
- Check that trajectory files do not exceed the current maximum size (10 GB as of June 2024).

If all of these checks pass, the simulation files are then copied from the *iRODS* landing directory to the task processing server. The following processing steps are then taken:

- Using MDTraj (39), save or convert the structure file to the “.pdb” format.
- Using MDTraj, save or convert the trajectory file to an “.xtc” file (which provides modest compression).
- Compute a hash of the topology file. If the topology hash matches other hashes already stored in the database, but the trajectory hashes do not, then the simulation is a replicate of an existing contribution, initialized from the same starting topology. The simulation is automatically linked with the existing contribution matching the hash.
- Compute RMSD and RMSF values of the trajectory.
- Produce a copy of the simulation, with extra atoms (lipids, water, and ions) removed from the simulation files using VMD (40)).
- Create a thumbnail image of the protein structure using the minimal structure file and NGL viewer (41).
- Extract the protein sequence from the structure file, and store it in the database entry for the simulation.
- Using MDTraj, create a down-sampled trajectory with 100 frames; this serves as the lightweight visual presented on the webpage for the simulation.
- Save database objects corresponding to the simulation, metadata, and simulation upload logs.
- Upload files from the simulation processing server to their final iRODS destination.

### Seeding MDRepo with simulations

Many repositories of MD simulations have been created over the years (e.g. (10–16)), each containing hundreds or thousands of simulations. Each of these repositories, typically containing simulations produced by the database hosts, is a valuable resource that provides researchers access to a quantity and diversity of simulations that they would be unlikely to produce on their own. However, data organization and access patterns differ between services, and relatively low server bandwidth means that batch downloads are generally quite slow. With the aim of improving accessibility of these data for researchers (relatively simple search/download protocols and improved access speed), we have imported simulations from two of these databases into *MDRepo*. The first data source is the ATLAS repository (16), which holds the results of simulations based on pdb-sourced topology files. At the time of download (April, 2024), the ATLAS website described the results as “freely available”, though no specific licence is described. Download of 2798 simulations from ATLAS required nearly three weeks to complete due to download bandwidth limitations. The second data set is GPCRmd (13), which was released under the same Creative Commons Attribution 4.0 license as *MDRepo*. At the time of download (January 2024), we retrieved 1457 simulations across 48 G-protein coupled receptor proteins, with download requiring over one week to complete. Each resulting *MDRepo* entry contains a reference (a “linkout”) to the URL associated with the simulation source at the time of import, to ensure that proper credit is given to data creators. (Note: since our download, the GPCRmd site has moved behind a registration wall, and many of the retrieved simulations seem to be unavailable on the site. This change in accessibility for data released under an open license highlights one of the motivations for an enduring and perpetually open simulation repository).

In addition to simulations gathered from two existing repositories, we have received several dozen contributions during an invitation-only phase of system validation. We intend to world with developers of other repositories to provide a secure and long-term storage option for their data, and we anticipate that the number of contributions from individual data creators will grow in the coming months.

## Discussion

MDRepo is an open repository for community-generated MD simulations of biomolecules. It is designed to provide a home for millions of simulations accumulated over years of research effort, with a robust storage infrastructure that ensures both data safety and high throughput data access. The centralized and open access nature of the repository will help to meet demands for reduced environmental impact by reducing redundant effort, improving reproducibility, and obviating dispersed storage solutions. Meanwhile, the anticipated 10s of petabytes of simulation data will enable new discoveries and modeling techniques.

A researcher may submit any number of simulations to MDRepo, from a single trajectory to thousands. Submitted simulations are subjected to some post-submission validation and preparation, then stored in infrastructure backed by CyVerse. Each simulation is stored as a separate entry, with standardized metadata captured for each. MDRepo places no restrictions on the use or distribution of stored data.

Users can search and explore simulations submitted by others. An individual trajectory can be downloaded directly from the website. Downloading a batch of simulations is performed with the *MDRepo* command-line tool.

We have seeded the repository with several thousand simulations, some gathered from other valuable repositories, and some newly generated for the repository. While this initial seed will serve as a large valuable resource to the community, it is only a first step; the promise of MDRepo will only be reached through extensive data contribution from the community. We anticipate that these data will enable new discoveries via re-analysis of individual simulations and through development of new Machine Learning models designed to leverage the rich trove of training data. We welcome the opportunity to work with the broader community to extend the collection of stored simulations into the millions, and to improve the functional and analytical features of the website. One important benefit of the architectural design of *MDRepo* is that the data are stored in a location that is designed to support access from academic and corporate cloud systems. Though the functionality does not yet exist, in the future we will establish the infrastructure to allow researchers to avoid the step of downloading data to their own servers, and instead to bring containerized analysis pipelines and machine learning models close to the data, for analysis in the cloud. While the current design ensures that data creators can be credited for their contributions to data found in *MDRepo* (through a combination of contributor lists, paper citations, and linkouts), we recognize the importance of improving the landscape of credit; over the coming months, we will formalize a robust framework for microcitation, so that researchers who contribute simulations to *MDRepo* will receive credit when those simulations are used by work leading to publications by other researchers.

## Acknowledgements

We thank members of the CompbioAsia research community, and the associated Bioinformatics Research Consortium, for discussions that initiated development of MDRepo, with particular gratitude for early suggestions by Charles Laughton and Chandra Verma. We also gratefully acknowledge the high performance computing (HPC) resources supported by the University of Arizona TRIF, UITS, and Research, Innovation, and Impact (RII) and maintained by the UArizona Research Technologies department.

## Funding

This work was made possible by support from the National Science Foundation under DBI Grant Nos. 0735191, 1265383, and 1743442, along with support from the University of Arizona Research, Innovation & Impact (RII) through BIO5 and IT4IR TRIF Funds. Computational workload depends on Jetstream2 at Indiana University through allocation BIO230080 from the NSF ACCESS program, which is supported by OAC Grants Nos. 2138259, 2138286, 2138307, 2137603, and 2138296. Author interaction and project conception was facilitated by NSF IRES grant 1953405.

## Competing interests

The authors declare no competing interests.

## Notes

https://mdrepo.org/

### Competing Interest Statement

The authors have declared no competing interest.

